# DNA Adenine Methylation Clock in Brain Aging and Alzheimer’s Disease Progression

**DOI:** 10.64898/2026.03.05.709867

**Authors:** Abdur Rahim, Xiaomei Zhan, Qiyuan Han, Allysha O’Donnell, Angela Jeong, Guru-Swamy Madugundu, Suresh Pujari, Monica Kruk, Xinlong Luo, Ling Li, Tao P. Wu, Natalia Y. Tretyakova

## Abstract

N^6^-methyldeoxyadenosine (N^6^medA) is a recently identified endogenous DNA modification widely found in bacteria, plants, and eukaryotes. In mammals, N^6^medA has been implicated in brain function, immunity, and response to environmental stress, but its relevance to gene regulation and mammalian aging remains controversial due to its extremely low abundance (< 1 per 10 million adenines) and an uncertainty regarding its genomic origin. We have developed and validated an ultrasensitive isotope dilution nano liquid chromatography-nanospray ionization Orbitrap mass spectrometry methodology to quantify N^6^medA in genomic DNA. Applying this approach to human prefrontal cortex tissues, we found that genomic N^6^medA levels increase linearly with chronological age (Pearson correlation coefficient, 0.95). Individuals with mild cognitive impairment (MCI) and Alzheimer’s disease (AD) exhibited a trend toward elevated cortical N^6^medA levels relative to age-matched controls. Genome-wide profiling of N^6^medA in human prefrontal cortex was conducted using two independent methods: NAME-Seq and MeDIP-Seq, which revealed age associated adenine methylation changes reminiscent of established epigenetic aging signatures such as the 5-methylcytosine clocks. N^6^medA mapping experiments identified a subset of genomic loci that were altered in MCI and AD. Pathway analysis of cross-validated adenine methylation sites revealed an enrichment of genes involved in neuronal function and age-related neurological processes, including glutamatergic synapse, axon guidance, and long-term depression. Finally, mass-spectrometry-based photoaffinity proteomics with synthetic DNA representing a region of the APP gene identified N^6^medA reader proteins with known roles in DNA repair, replication and transcription. Together, these findings identify N^6^medA as an age-associated DNA modification in the human brain and suggest that its accumulation and recognition by specific protein readers may contribute to molecular processes underlying brain aging and age-related neurodegeneration.

## Introduction

Aging is a major risk factor for Alzheimer’s disease and other dementias, which manifest as a loss of synaptic plasticity, memory impairment, and defective learning capabilities.^1,2^ Alzheimer’s disease (AD) is the most common type of dementia (60%-80%), affecting around 6.5 million individuals in the United States.^1^ Accumulation of the β-amyloid protein, which leads to amyloid-beta plaques, tau tangles, synapse loss, inflammation, and neurodegeneration, is generally considered the primary initiating event in Alzheimer’s disease (the amyloid hypothesis).^3^ While the number of AD cases is expected to double by the year 2060,^1^ AD treatment strategies remain limited, with nearly 99% of clinical trials facing failure and about 200 drug development programs abandoned in the past decade.^4,5^ With a rare exception of recently approved anti-amyloid antibodies,^6^ drugs targeting amyloid-beta plaques in the brain have failed, highlighting an urgent need for new therapeutic targets for AD.

Age-related brain deterioration and neurodegeneration pathologies are often characterized by epigenetic deregulation.^7^ Specifically, changes in DNA methylation mark 5-methylcytosine (5mC), are well known to be associated with brain aging in humans. 5mC represents 3-5% of total Cs and is a key epigenetic mark that controls the levels of gene expression by mediating transcription factor binding and chromatin remodeling.^8^ Aging is characterized by global cytosine hypomethylation, along with changes in methylation patterns of specific genomic loci, leading to altered chromatin structure.^9^ This has led to the development of “epigenetic clocks”, which estimate biological age by measuring DNA methylation levels.^10–12^ Cytosine methylation patterns have been shown to be valuable biological age prediction tools, able to forecast the life span with accuracy surpassing the models based on telomere length.^13^ Additionally, cytosine methylation is associated with in brain aging and the development of AD. De Jager et al. mapped cytosine methylation and gene expression patterns in >700 autopsied human brain samples, revealing significant cytosine methylation changes in AD patients relative to controls, these included *ANK1*, *CDH23*, *DIP2A*, *RHBDF2*, *RPL13*, *SERPINF1*, and *SERPINF2* genes.^14^ The neuropathological markers, such as neurotic plaques, diffuse plaques, and amyloid load, were associated with epigenetic age acceleration in the dorsolateral prefrontal cortex.^15^ However, there is no consensus on the mechanisms by which DNA methylation contributes to AD pathology, and it is not known whether additional epigenetic marks contribute to brain aging and AD etiology.

Genomic *N^6^*-methyldeoxyadenosine(N^6^medA) has long been known to play a role in modification-restriction systems in prokaryotes, but was only recently identified in metazoans. In 2015, three separate reports described the observation of N^6^medA in eukaryotic genomic DNA of *Chlamydomonas reinhardtii*, *Caenorhabditis elegans*, and *Drosophila melanogaster*.^2,16,17^ The reported levels of N^6^medA were around 0.4 % of adenines in *Chlamydomonas*,^2^ ranged from 0.013% to 0.39% of adenines in *C. elegans*,^16^ and ∼0.001% to 0.07% in *Drosophila*.^17^ In *Chlamydomonas*, N^6^medA was mainly located at the ApT dinucleotides around transcription start sites (TSS) and showed a bimodal distribution, localizing primarily within the linker DNA between nucleosomes and marking actively expressed genes.^2^ In *C. elegans,* a broad distribution of N^6^medA across all chromosomes was identified, with no significant enrichment or depletion across genomic features.^16^ In *Drosophila,* N^6^medA was proposed to promote the activity of transposons.^17^ Charria et al. reported that N^6^medA was widespread across genomes of diverse eukaryotes and was preferentially localized near active genes.^18^ Wu et al. have shown that N^6^-adenine DNA methylation occurs in mammalian embryonic stem cells (ESCs), where it acts as a repressive mark that silences transcription of long interspersed nuclear element (LINE) transposons.^19^ N^6^medA has been reported in genomic DNA isolated human tissues based on the results of single-molecule real-time sequencing, but these results have been later questioned due to its low abundance and a high probability of false positives.^20,21^ [G/C]AGG[C/T] has been recognized as the sequence motif most significantly associated with N^6^medA formation.^19,20^ Further, it was reported that N^6^-adenine methylation is enriched in gene coding regions and marked actively transcribed genes in human blood cells.^20^

Much controversy exists regarding N^6^medA writers and readers. Initially, N^6^-methyladenine transferase (N^6^AMT1) was identified as an N^6^medA writer and ALKBH1 as an N^6^medA eraser in mammalian cells.^19,20^ However, other studies questioned the ability of N^6^AMT1 to generate N^6^medA in DNA^20,22^ and dismissed the role of N^6^AMT1 in DNA methylation in humans.^18^ Xie at al. have shown that N^6^medA levels were dynamically regulated by demethylase ALKBH1, and that small molecule inhibitors of ALKBH1 reduced glioblastoma cell proliferation.^23^ Collectively, these findings led to a hypothesis that N^6^medA plays a role in epigenetic regulation.

N^6^medA has been reported as an epigenetic indicator of early life stress,^24^ a dynamic mark affected by environmental stress in a mouse model,^25^ and as a DNA modification playing a regulatory role in the formation of fear extinction memories.^26^ A recent report by Fernandes et al. implicated N^6^medA in embryogenesis and fetal development and demonstrated that tissue levels of N^6^medA steadily increase with age in mice and zebrafish.^27^ Similarly, Sturm et al reported that mitochondrial levels of N^6^medA increases with age in *C. elegans*, Drosophila, and dogs.^28^ A study by Xie at al. revealed that N^6^medA is markedly increased in glioblastoma and co-localized with heterochromatic histone modifications, predominantly H3K9me3.^23^

The extremely low genomic abundance of N^6^medA and the high probability of false positives in its detection have led to skepticism regarding its possible role in epigenetic regulation.^29^ Musheev et al. proposed that methylated dA is produced via metabolism of N^6^-methyladenosine (N^6^meA) present in the nucleoside poll, followed by fortuitous incorporation of N^6^-Me-dAMP into DNA during DNA replication.^30^ In addition, large variations in global N^6^medA quantification ranging from 0.051% of total As to 1 ppm^31^ led to speculations that global N^6^medA in mammalian systems was overestimated due to possible bacterial contamination, RNA contamination, and technical limitations of the methods used to quantify this mark.^32,33^

In the present study, we developed an ultrasensitive and artifact free isotope labeling methodology for genomic N^6^medA utilizing Orbitrap nanoLC-NSI-MS/MS. Using this new approach, we accurately quantified global N^6^medA levels in DNA isolated from prefrontal cortex of humans 15-95 years of age and in age matched individuals with no cognitive issues, mild cognitive impairment, and Alzheimer’s disease. Additionally, we mapped N^6^medA across the human genome using two independent methodologies (NAME-Seq and MeDIP-Seq) and characterized protein readers of this novel epigenetic modification within the APP gene using mass spectrometry based photoaffinity proteomics.

## Results

### Quantification of N^6^medA in genomic DNA isolated from prefrontal cortex: effects of age and AD development

To enable accurate quantification of global N^6^medA in genomic DNA, any possibility for bacterial contamination must be eliminated, and the method must be sensitive enough to detect rare epigenetic marks. We chose Orbitrap nanoLC-NSI-MS/MS due to it high mass accuracy and sensitivity in analyzing rare DNA modifications.^34^ Sample preparation procedures including DNA extraction, enzymatic digestion, and solid phase extraction (SPE) were carefully optimized to minimize any contamination introduced during sample preparation. Nano HPLC interfaced with an Orbitrap QE mass spectrometer was used to allow for high resolution mass measurements required for trace level analysis of DNA modification. Isotopically labeled N^6^medA was synthesized in our laboratory and used as an internal standard for mass spectrometry. Isotope dilution nanoLC-ESI^+^-MS/MS methodology was validated by analyzing samples prepared by spiking known amounts of N^6^medA-containing synthetic DNA oligodeoxynucleotides into ^15^N-labeled genomic DNA from *E. coli* that was cultured with ^15^NH_4_Cl. To control for any possible contamination, each analysis included a negative control containing all enzymes, solvents, and an internal standard in the absence of genomic DNA. With careful optimization, we developed a method that can accurately quantify N^6^medA in DNA within a range from 0.05 fmol/µg DNA to 3 fmol/µg DNA (**Figure 1A**), with a clean or negligible background in our sample preparation control (**Figure 1B**).

**Fig. 1:**
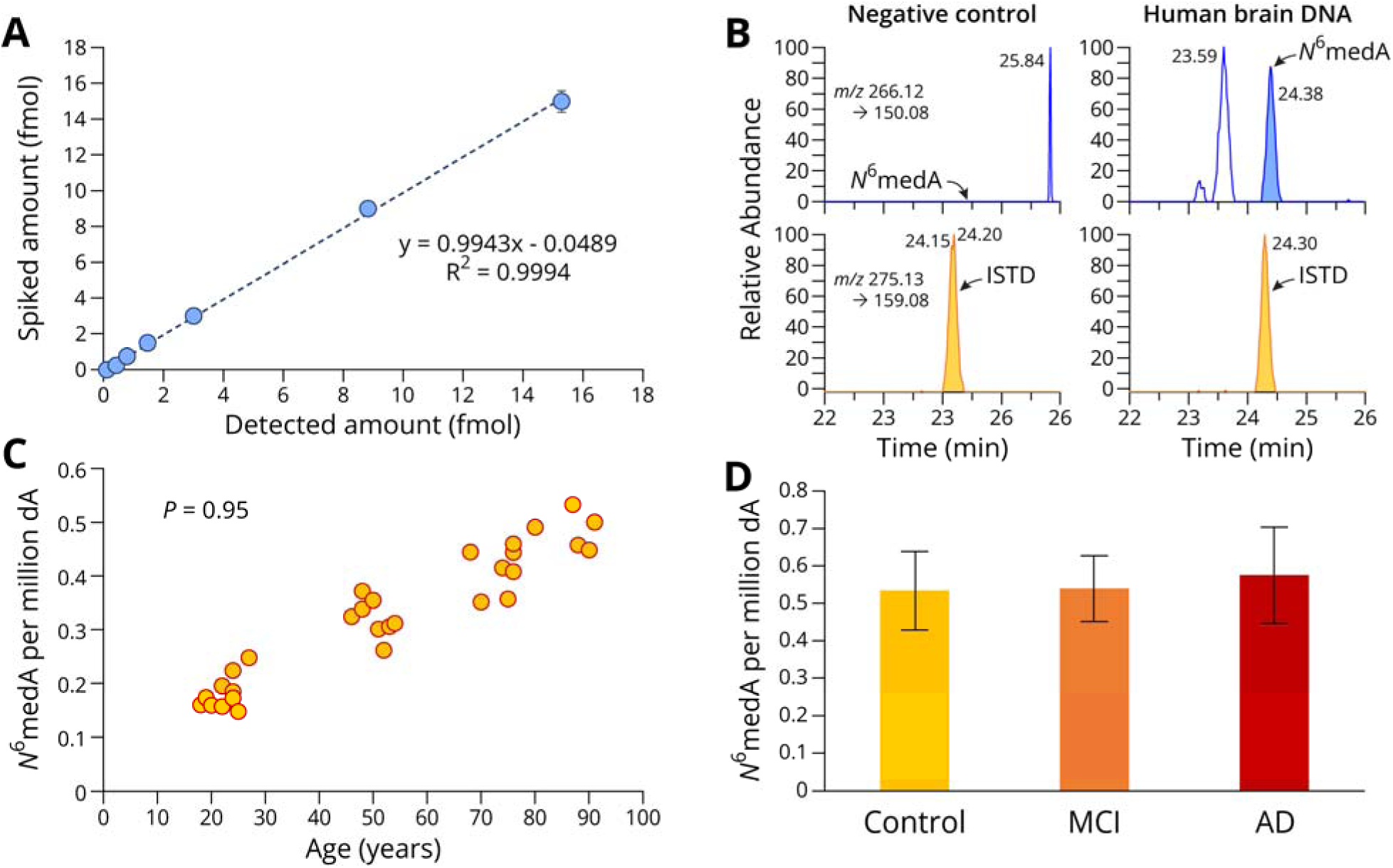
NanoHPLC-NSI-HRMS/MS reveals age- and AD-associated differences in global N^6^medA levels in genomic DNA isolated from human prefrontal cortex. **A:** NanoHPLC-NSI-HRMS/MS method validation for N^6^medA. ^15^N-labeled genomic DNA was spiked with increasing amounts of N^6^medA-containing synthetic oligodeoxynucleotides, followed by enzymatic digestion to deoxynucleosides, SPE, and nanoLC-NSI-MS/MS analyses on Oribitrap QE MS. **B:** Representative nanoHPLC-NSI-HRMS/MS traces for N^6^medA in a negative control sample (left) and in a human brain DNA sample (right). **C,** Scatter plot showing a significant positive correlation between chronological age and global N^6^medA level in genomic DNA isolated from the prefrontal cortex region of healthy human brains. **D,** Global N^6^medA levels in the prefrontal cortex region of human subjects: healthy subjects (CNTRL), individuals with mild cognitive impairment (MCI) and AD patients (AD). Each group contained 14-16 individuals aged between 87.8 and 90 years old (*Table 1*).

**Table 1:** Human participant demographics for data shown in Figure 1D.

|  | **NCI (No cognitive impairment)** | **MCI (Mild cognitive impairment)** | **AD (Alzheimer's dementia)** |
| --- | --- | --- | --- |
| # Samples | 16 | 14 | 15 |
| Age at death (yr) | 89.3 ± 4.1 | 88.7 ± 6.8 | 90.1 ± 3.9 |
| Male/female | 6/10 | 7/7 | 6/9 |
| Education (yr) | 18.1 ± 2.6 | 17.4 ± 3.9 | 18.5 ± 3.2 |
| Post-mortem interval (hr) | 5.99 ± 5.48 | 6.25 ± 3.34 | 5.59 ± 3.07 |
| Total amyloid-β (% area) | 0.75 ± 0.99 | 1.07 ± 1.74 | 6.23 ± 2.78 |
| Tau tangle density | 1.49 ± 1.15 | 2.34 ± 1.86 | 9.64 ± 6.26 |

The validated Orbitrap nanoLC-NSI-MS/MS methodology was applied to quantify global N^6^medA levels in genomic DNA isolated from prefrontal cortex region in autopsied human brain samples. Prefrontal cortex region of the brain was selected due to its central role in cognitive control and executive functions, as well as its involvement in AD pathology.^35,36^ Among the human aging cohort samples, which included healthy brains at different ages, we observed a strong positive correlation between global N^6^medA levels (0.15-0.53 N^6^medA per million dA) and subject’s chronological age, with a Pearson correlation coefficient of 0.95 (**Figure 1C**). To the best of our knowledge, this is the first example of showing an increase of genomic N^6^medA levels in human brain with age.

We further quantified N^6^medA in genomic DNA extracted from the prefrontal cortex of healthy seniors aged between 83.4 and 96.1 years old, patients with diagnosed MCI aged between 76.9 and 97.6 years old, and AD patients aged between 82.8 and 98 years old (**Table 1**). NanoLC-NSI-MS/MS analyses revealed that global N^6^medA levels increased in DNA of MCI patients (0.53 ± 0.086 N^6^medA per million dA), and AD patients (0.57 ± 0.127 N^6^medA per million dA), as compared to age matched healthy controls (0.53 ± 0.104 N^6^medA per million dA). These differences were not statistically significant (**Figure 1D**), probably due to limited number of samples (14-16 per group).

### Mapping N^6^medA distribution across the genome by MeDIP-seq

In the event that N^6^medA serves as an epigenetic mark in the human brain, it should be distributed non-randomly across the genome. To address this question, adenine methylation marks were mapped across the genome of a healthy 19-year old female, a healthy 81.8-year old female, and a female AD patient 82.5 years of age (**Table 2**). Genome wide mapping of adenine methylation marks was initially conducted by N^6^medA immunoprecipitation in combination with next generation sequencing (MeDIP-seq).^25^

**Table 2:** Sample information in the study of genome wide mapping of N^6^medA in human brains.

| Sample ID | Sex | Age | Number of N<sup>6</sup>medA peaks |
| --- | --- | --- | --- |
| CNTR 19 Y.O. | Female | 19 | 356136 |
| CNTR 81.8 Y.O. | Female | 81.8 | 287679 |
| AD 82.5 Y.O. | Female | 82.5 | 356526 |

The number of N^6^medA peaks measured by N^6^medA IP-seq in human prefrontal cortex ranged from 287,679 to 356,526 per sample, with both common and unique peaks observed in each subject (**Table 2**, **Figure 2A**). N^6^medA peak count frequency across the genome indicated an enrichment at gene body and distal intergenic regions, but was relatively depleted at transcription start and transcription termination sites (**Figure 2B**). Genes containing unique N^6^medA peaks in the AD patient as compared to matched healthy control of similar age were enriched in KEGG pathways such as GABAergic synapses, glutamatergic synapses, cardiomyopathy, and serotonergic synapses (**Figure 2C**). This may be relevant to AD etiology as correct functioning of glutamatergic synapses is essential for memory, emotional regulation, and learning.^37^ Genes containing unique N^6^medA sites in genomic DNA of senior healthy female as compared to the AD patient were enriched in KEGG pathways corresponding to lactin regulation, autophagy, and yersinia infection (**Figure 2D**). N^6^medA peaks in the brain of the senior healthy female sample as compared to the young healthy control were enriched in the ErbB signaling, inositol phosphate metabolism, and adherens junction pathways (**Figure 2E**). Collectively, our **MeDIP-seq** results suggest that N^6^medA is distributed non-randomly across the genome and that DNA adenine methylation patterns are influenced by donor’s age and Alzheimer’s disease pathology.

**Fig. 2:**
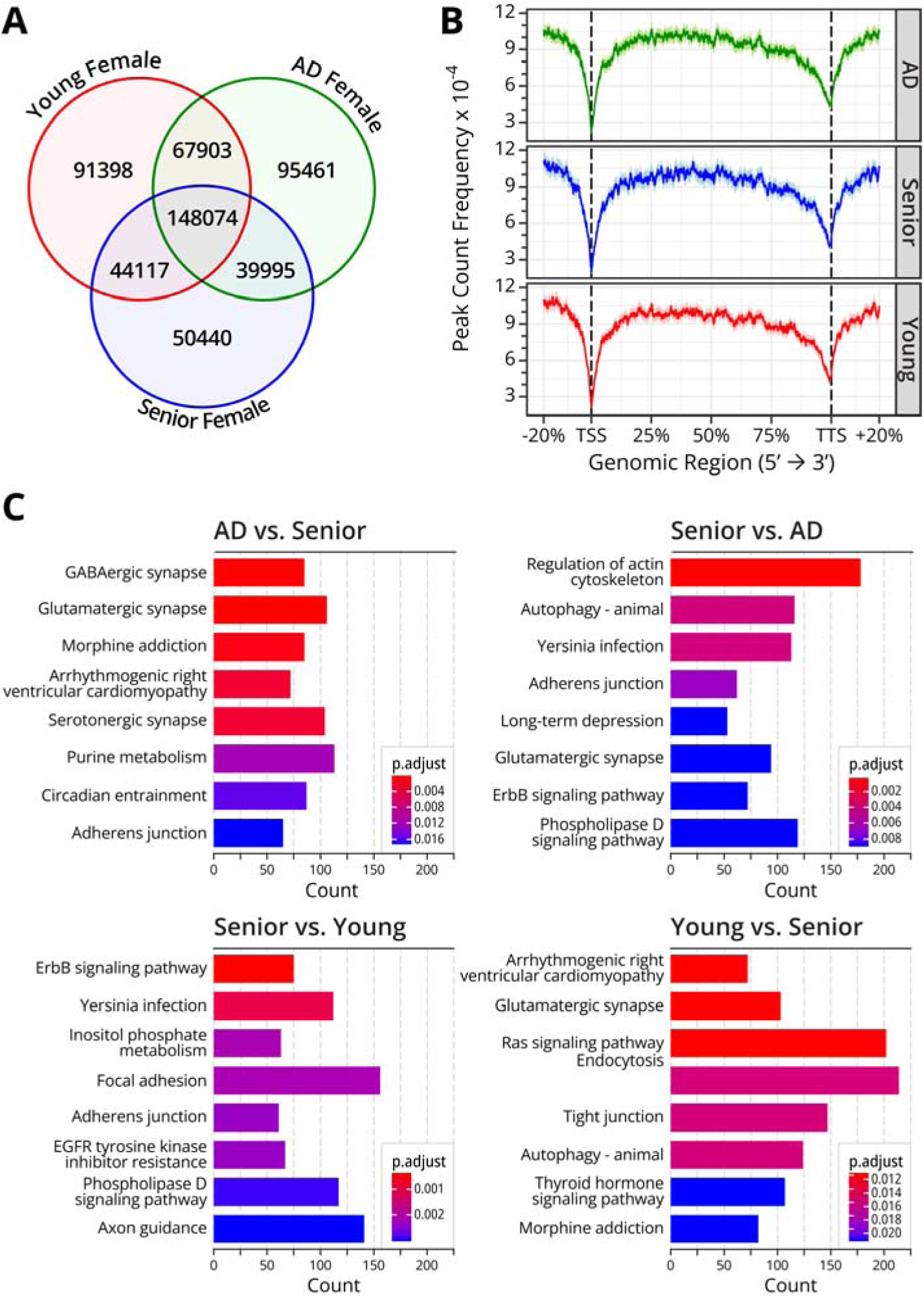
MeDIP-seq results for N^6^medA mapping in genomic DNA isolated from prefrontal cortex of 3 human subjects: a young female (19 y.o.), a senior female control (81.8 y.o). and a female AD patient (82.5 y.o). **A,** Venn diagram showing unique and shared genomic N^6^medA peaks found in the prefrontal cortex of a young female (19 y.o.), aged female control (81.8 y.o). and a female AD patient (82.5 y.o). **B,** N^6^medA peak frequency across genomic regions. **C,** KEGG pathway analysis of genes containing unique N^6^medA peaks in AD patient and aged matched control when comparing these two groups and senior and young when comparing these two groups.

### Mapping N^6^medA across human genome by NAME-seq

Since MeDIP-seq. methodologies are known to have high false positive rates due to antibody cross-reactivity, low signal to noise ratios, and possible library preparation bias,^38–40^ N^6^-me-dA profiling in the same subjects was repeated using <u>N</u>itrite-assisted <u>A</u>mino <u>ME</u>thylation sequencing (NAME-seq), an enzyme-based N^6^medA-to-T mutation sequencing method.^41^ In NAME-seq, genomic DNA is treated with nitrite, converting A to I and 6mA to nitrosylated-6mA (6mA-NO). This conversion is followed by Klenow fragment (3’→5’ exo^−^) mediated DNA synthesis to specifically induce the 6mA-to-T transversion mutations at the N^6^medA sites.^41^

NAME-seq methodology was used to map adenine methylation marks across the cortical genome of a healthy female aged 19 (CNTRL 19 y.o.), a healthy female aged 81.8 (CNTRL 81.8 y.o.), a female mild cognitive impairment patient aged (MCI 82.9 Y.O.), and a female AD patient aged 82.5 years (AD 82.5 Y.O.) (**Table 3**). The number of N^6^medA peaks measured by NAME-seq in human prefrontal cortex ranged from 26,628 to 53,198 per sample, with both shared and unique peaks observed in each case (**Table 3** and **Figure 3A**). Overall, the number of N^6^medA peaks detected by NAME-seq (**Table 3**, **Figure 3**) was approximately 10-fold lower than MeDIP-seq (**Table 2**, **Figure 2A**), consistent with high false positive rates of the latter. KEGG pathway analysis of genes containing N^6^medA modification revealed an enrichment of genes involved in focal adhesion, glutamatergic synapse, axon guidance, and morphine addition pathways (**Figure 3C)**. As mentioned above, glutamatergic synapses play a vital role in emotion regulation, learning and memory,^37^ while axon guidance is known to have roles in synapse formation and other key neurotransmission processes in the brain.^42,43^

**Fig. 3:**
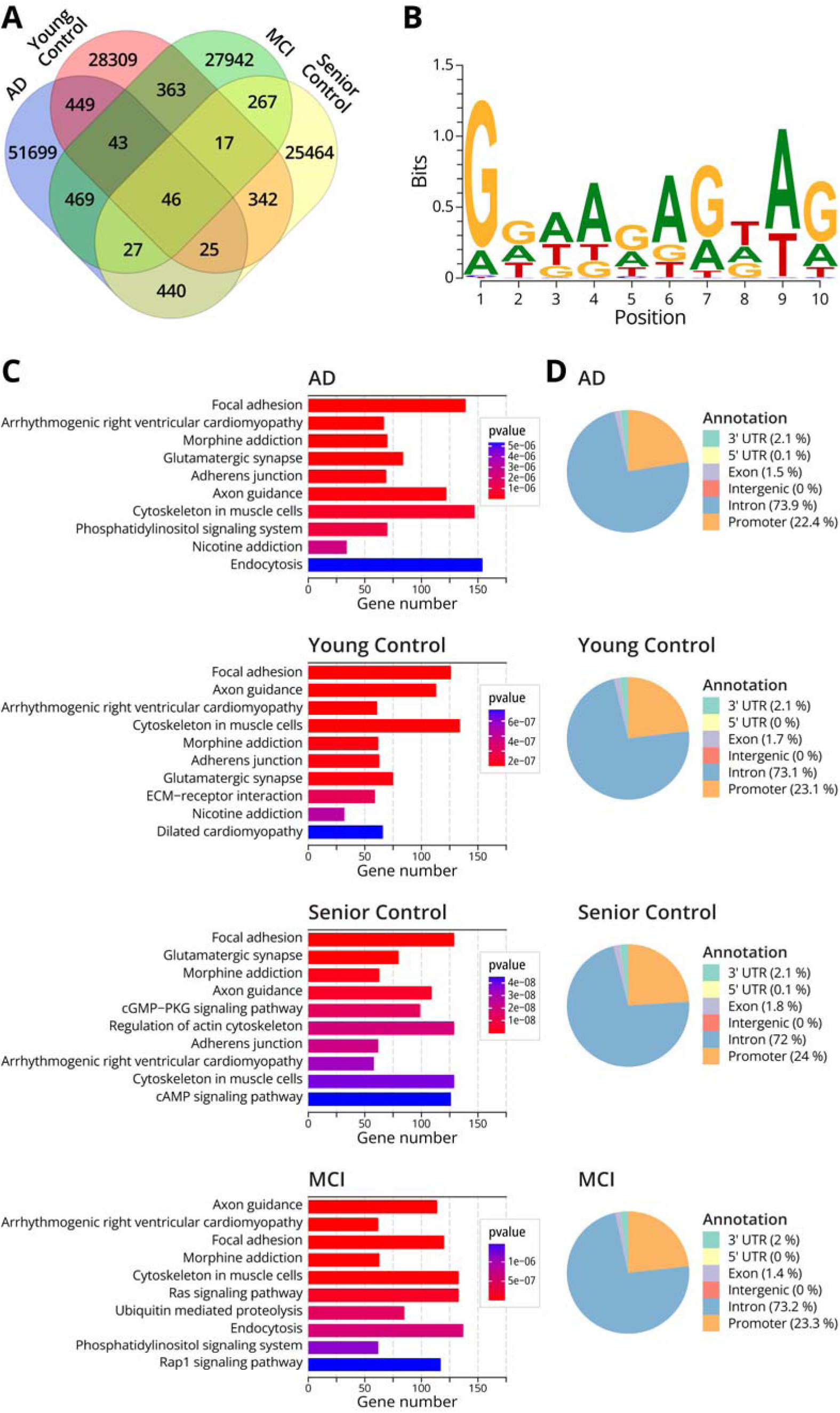
NAME-seq results for N^6^medA mapped in genomic DNA isolated from prefrontal cortex of four human subjects: a young female (19 y.o.), a senior female control (81.8 y.o), a female with mild cognitive impairment (MCI, 82.9 y.o.), and a female AD patient (82.5 y.o). **A,** Venn diagram showing the number of N^6^medA peaks identified in the prefrontal cortex of four individuals. **B,** Top sequence motif for N6-adenine methylation in AD female patient. **C,** KEGG pathway analysis of genes containing N^6^medA peaks in AD female, senior healthy female and young healthy female. **D,** Gene annotation analysis of genes containing N^6^medA peaks in a female subject with AD, a female with MCI, an age matched senior healthy female, and a young healthy female.

**Table 3:** Sample information in the study of genome wide mapping of N^6^medA in human prefrontal cortex using NAME-seq methodology.

| Sample ID | Sex | Age | N<sup>6</sup>medA peaks |
| --- | --- | --- | --- |
| CNTR 19 Y.O. | Female | 19 | 29594 |
| CNTR 81.8 Y.O. | Female | 81.8 | 26628 |
| MCI 82.9 Y.O. | Female | 82.9 | 29174 |
| AD 82.5 Y.O. | Female | 82.5 | 53198 |

Next, we carried out a GO enrichment analysis on genes containing N^6^medA peaks in all four samples in an attempt to interpret the function of adenine methylation in three main categories: biological processes (BP), molecular functions (MF), and cellular components (CC). Important BP such as dendrite development and synapse organization were observed for genes associated with adenine methylation (**Supplementary Figure S1a-d**). Similarly, genetic functions in cellular components such as glutamatergic synapse and synaptic membrane were also identified (**Supplementary Figure S1a-d).** Taken together, our NAME-Seq results suggest a potential role of N^6^medA in learning and memory.

Genomic annotation analysis of the N^6^medA peaks revealed that among various elements, intron regions of genes had the highest probability of N^6^medA peaks, followed by promoters, 3’-UTR, and exons (**Figures 3D**). Further, we identified sequence motifs for N^6^-adenine methylation in the human brain. For AD and MCI samples, N^6^medA was preferentially found in the GGAA motif, while for the in healthy brain sample, adenine methylation was observed preferentially in the CCCA motif (**Figure 3B, Supplementary Figure S1e-f**).

### Integration of N^6^medA NAME-seq. and Me-DIP seq results

Although the N^6^medA MeDIP-Seq methodology has been previously used for adenine methylation profiling, the sensitivity and specificity of this approach have been debated.^23,40,44^ A major concern for MeDIP-Seq analyses of N^6^medA is its high false positive rates, especially given the low overall abundance of the adenine methylation marks.^40^ Indeed, our NAME-seq analysis of human brain DNA samples yielded 10-fold fewer N^6^medA peaks than did MeDIP-Seq (**Tables 2, 3**). To cross-validate our results, additional bioinformatics analyses were conducted to identify N^6^medA sites that were independently detected by both MeDIP-seq and NAME-seq methodologies. Approximately 10% of N^6^medA sites identified in the NAME-seq experiments were cross-validated by the MeDIP-Seq methodology, which is significantly higher than the random background (**Table 4, Supplementary Figures S2a-c**). KEGG pathway analysis of the validated sites indicated that N^6^medA is overrepresented in genes involved in glutamatergic synapses, axon guidance, long-term depression, and several other pathways (**Figure 4A**). Subsequently, we carried out a GO enrichment analysis on the overlapped genes to interpret genes function in three main categories such as biological process (BP), molecular function (MF), and cellular component (CC). Genomic adenine methylation was associated with genes involved in axon guidance, oxytocin signaling, glutamatergic synapse, long term depression, and morphine addiction. Accurate axonogenesis is required for axon guidance and synapse formation,^45^ while synapse organization has a role in neuronal communications.^46^ As discussed above, similar pathways were identified upon analyses of IP-seq and NAME-seq data (**Figures 2**–**3, Supplementary Figure S3**). Consistent with our nanoLV-NSI-MS/MS data (**Figure 1**), the AD patient brain sample contained a greater number of adenine methylation sites (**Tables 3,4**).

**Figure 4:**
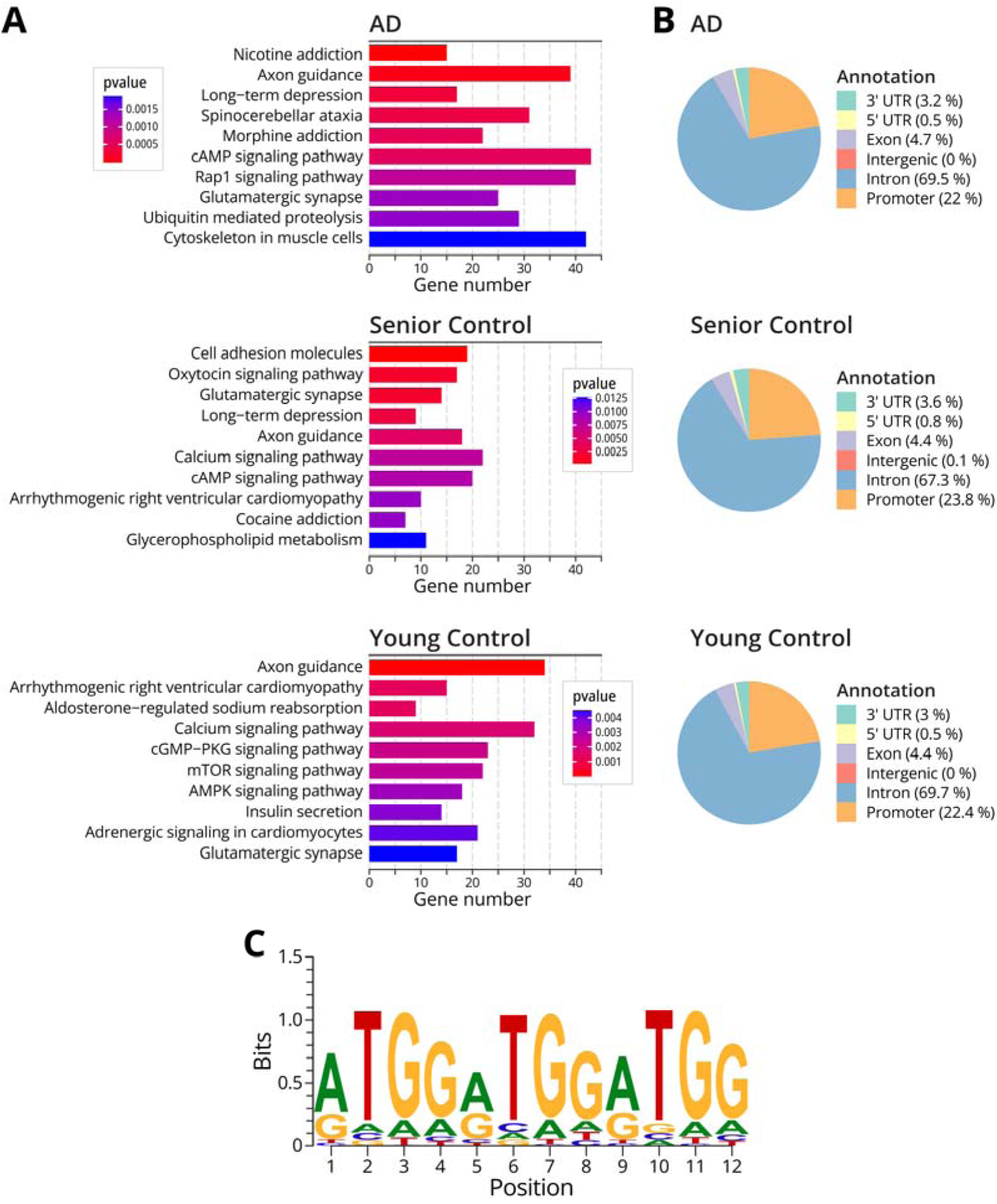
KEGG pathway enrichment analysis, gene annotation, and motif discovery for NAME-seq and IP seq. overlapped sites. **A,** KEGG pathway analysis of the cross-validated genomic N^6^medA sites detected in the prefrontal cortex of a female AD patient, a senior healthy female, and a young healthy female, respectively. **B,** Genomic annotation of the overlapped sites for AD patient, senior healthy female, and young healthy female respectively. **C,** Top sequence motif for the N^6^medA sites confirmed by both methods in the AD patient sample.

**Table 4:**
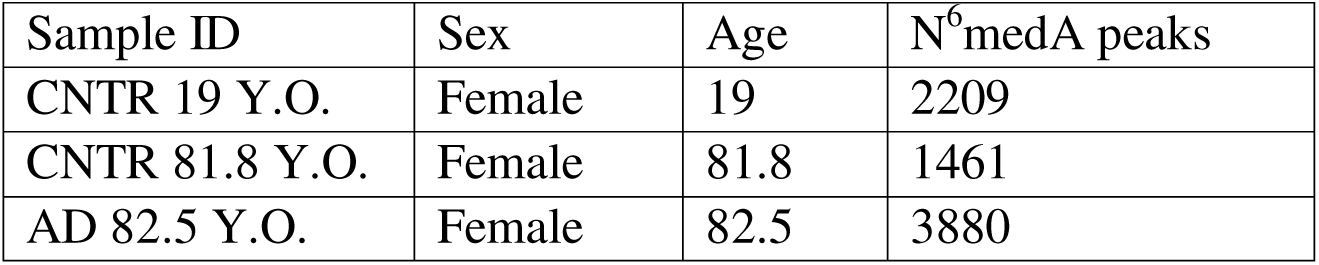
Total numbers of cross-validated N^6^medA peaks detected using both MeDIP-seq and NAME-seq methodologies in genomic DNA isolated from prefrontal cortex of 3 human subjects: a female AD patient (82.5 y.o), a senior female control (81.8 y.o), and a young female (19 y.o.).

Genomic distribution analysis of the cross-validated sites revealed that N^6^medA loci were enriched in the intron regions, followed by promoters, exons, and 3’-UTR (**Figure 4B**). Sequence motif searching indicated that the “ATGG” motif was overrepresented in the AD sample (**Figure 4C**). This is consistent with previous studies that reported that N^6^medA was enriched at AT -rich regions of mammalian genomes.^19^ However, for 81.8 y.o. healthy female, adenine methylation was observed preferentially in the CATC motif (**Supplementary Figure S3d**). Additional sequencing experiments and validation are needed to confirm these results.

### Affinity proteomics for specific protein readers in N^6^medA

A major mechanism by which DNA epigenetic marks exert their biological function is through recruiting specific protein “readers”.^47^ Previous studies have identified several potential N^6^medA readers including SSBP1, YTHDF3, and Histone 2AX.^48,49^ To characterize specific protein readers of N^6^medA in the brain, we conducted a photoaffinity proteomics experiments employing protein extracts from human neuroblastoma cell line SH-SY5Y and a N^6^medA containing DNA “bait” derived from the amyloid precursor protein (APP) gene. SH-SY5Y cells were selected because they are commonly employed in neuroscience research as an *in vitro* model for AD.^50^ The sequence of the DNA oligodeoxynucleotide was derived from a N^6^medA peak in the APP gene that contains the most enriched motif of N^6^medA according to our sequencing data see above). This sequence motif (CAGGC) is consistent with previously identified N^6^medA motif in human genome determined via SMART seq.^20^

In our affinity proteomics experiments, DNA duplexes containing a 5’-biotin tag, a centrally positioned dA or N^6^medA, and a diazirine photoaffinity group were incubated with nuclear protein extracts from SH-SY5Ycells. Following UV irradiation to cross-link protein readers to DNA, the resulting DNA-protein conjugates were purified via streptavidin beads and subjected to on bead tryptic digestion and nanoLC-NSI-MS/MS to identify and quantify the captured proteins (**Table 5**, **Figure 5A**). Affinity capture experiments were conducted in triplicate, and the MS based proteomics results were compared for N^6^medA containing DNA “baits” and the unmethylated dA controls. After employing student t-test at an FDR cutoff at 0.01, we identified 1355 proteins which were significantly enriched in N^6^medA pulldowns as compared to dA controls. These proteins are enriched in pathways such as the spliceosome, mRNA processing, nucleocytoplasmic transport, DNA repair, and DNA replication (**Figures 5B-D**). PRKRIP1, CAMSAP3, and ZRSR2P1 were some of the most significantly enriched proteins preferentially binding to N^6^medA DNA as compared to dA control (**Figure 5B**). PRKRIP1 (Protein Kinase R-Interacting Protein 1) is a component of the spliceosome that has a function in pre-mRNA splicing through double stranded RNA binding.^51^ RNA N^6^meA is widely documented to function in splicing, translation, and RNA stability,^52^ thus it is not surprising that our data provides evidence for an involvement of its DNA analog in these processes.

**Fig. 5:**
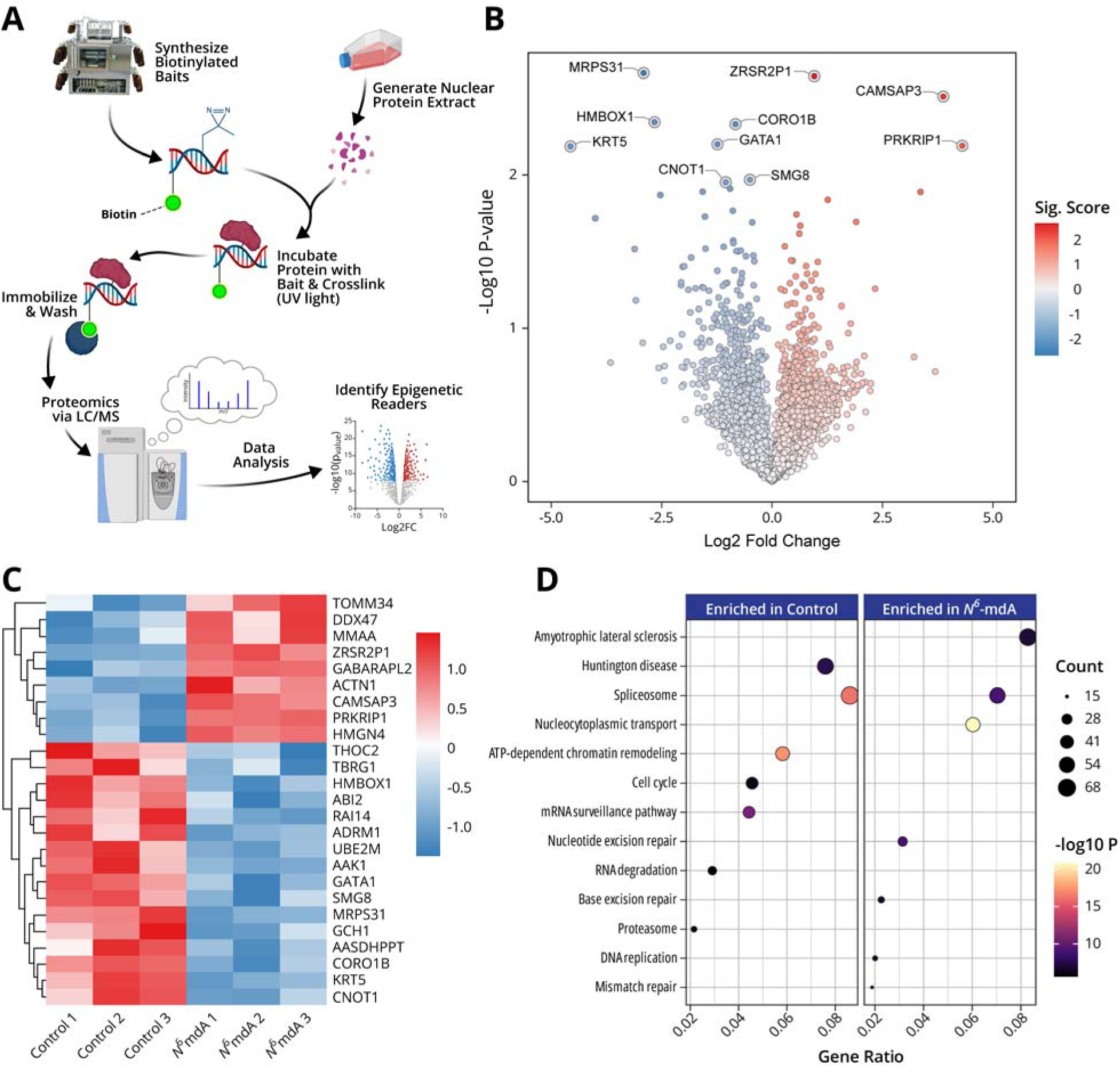
Characterization of protein readers of N^6^medA via photoaffinity proteomics. **A,** Schematic representation of the photoaffinity proteomics experiment to identify N^6^medA. **B,** Volcano plot quantitative analysis of proteins identified upon N^6^medA affinity pulldown in SH-SY5Y cells nuclear extract. **C,** Hierarchical clustering heatmap showing top 25 proteins preferentially identified upon N^6^medA photoaffinity pulldown in SH-SY5Y cells nuclear extract as compared to dA. **D,** KEGG pathway enrichment of proteins with preferential affinity for N^6^medA.

**Table 5:**
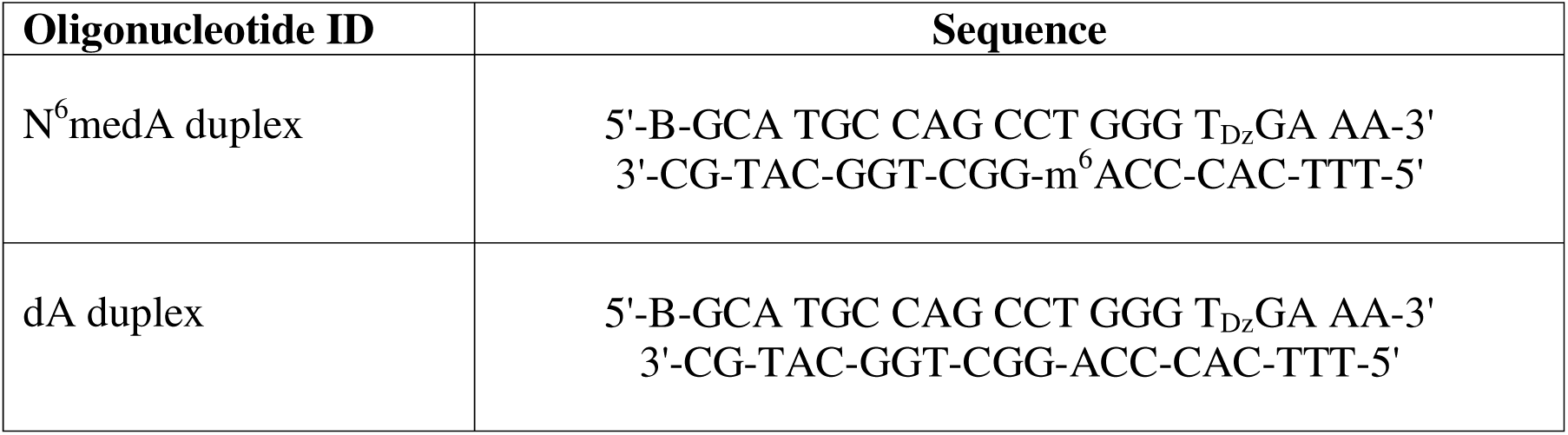
Nucleobase sequence of double stranded DNA used in photo-affinity proteomics assay for characterizing N^6^medA specific protein readers in the brain. Genomic coordination of this sequence is ch21:25969802-25969821 in human GRChg38. B: biotin, Dz: diazirine.

When the data was limited to proteins most significantly enriched upon N^6^medA capture (over 30%), we identified 25 proteins with significantly higher affinity for DNA N^6^medA as compared to dA (**Figure 5C**). These potential readers are known to function in DNA replication, transcription, and chromatin architecture. For instance, HMGN4 (high mobility group nucleosome-binding protein 4) plays a key role in chromatin remodeling by directly binding to nucleosomes, thereby influencing transcription, replication and repair.^53^ On the other hand, DDX47 (DEAD-box helicase 47) is a known a RNA helicase known to be involved in pre-rRNA processing, ribosome assembly, and nucleolar organization.^54,55^

Next, we repeated proteomics data analysis focusing specifically on gene products involved in DNA repair pathways. As shown in **Supplementary Figure S4,** several DNA repair proteins such as TRIM28, RPA1, RPA2, CHAF1A etc. were found to have an increased affinity for N^6^medA containing DNA as compared to dA. For example, TRIM28 is a nuclear scaffold protein that acts as a transcriptional repressor, epigenetic regulator, and a participant in DNA damage response and thereby maintain chromatin organization and genome stability.^56,57^ RPA1 (Replication protein A1) is the largest subunit of the RPA complex that binds and stabilizes single stranded DNA during replication and repair to coordinate DNA synthesis, damage signaling, and genome stability.^58,59^ CHAF1A is the p150 subunit of the chromatin assembly factor-1 (CAF-1) complex and functions as a histone chaperone that mediates replication-coupled nucleosome assembly by depositing newly synthesized histone H3–H4 onto nascent DNA through its interaction with PCNA.^60,61^ By coordinating chromatin reassembly during DNA replication and DNA repair, CHAF1A is essential for the maintenance of genome stability and epigenetic inheritance.^62,63^ Taken together, in the context of CX-GGC, the identification of these specific protein readers such as HMGN4, TRIM24, RPA1, CHAF1A suggests a possible role of DNA N^6^meA in regulating DNA repair, replication, and transcription.

## Discussion

In the present study, we examined genomic N^6^-adenine methylation in the human brain and evaluated its potential association with aging and Alzheimer’s Disease. A highly sensitive, fully validated nanoLC-NSI-MS/MS methodology was used to accurately quantify global genomic levels of N^6^medA in the prefrontal cortex of healthy controls of increasing age and in patients with MCI and AD (**Figure 1**). Further, genome-wide N^6^medA mapping in human brain DNA was carried out by two independent methodologies, affinity based MeDIP-Seq (**Figure 2**) and a novel chemical method, NAME-seq (**Figure 3, 4**).^41^ Finally, photoaffinity-based proteomics experiments revealed potential protein readers and “anti-readers” of adenine methylation marks in the context of the APP gene (**Figure 5**).

Our isotope labeling nanoLC-ESI-MS/MS results indicate that N^6^medA is a rare DNA mark in human brain (2-6 N^6^medA per 10^7^ adenines or 5,700 per cell) (**Figure 1**). This value is lower than previously published data for human samples,^31^ which could be explained by rigorous control of contamination in our study and/or by differences between tissue types. Our nanoLC-ESI-MS/MS results for N^6^medA (2-6 N^6^medA per 10^6^ As or 5,700 N^6^medA sites per cell) agree with the sequencing data for the same tissue. Specifically, our NAME-Seq experiments identified approximately 26,000-53,000 N^6^medA sites per cell (**Table 3**), while MeDIP-Seq detected 287,679-356,526 sites (**Table 2**), and the number of sites detected by both methods was approximately 3,000/cell (**Supplementary Table S1**). Thus, the number of sites confirmed by both methods (3,000 per cell) roughly matches our nanoLC-ESI-MS/MS data (5,700 per cell).

To the best of our knowledge, our study is the first to report that genomic N^6^medA levels in human brain increase with age (**Figure 1A**). However, N^6^medA abundance was previously reported to increase in tissues of zebrafish and mice across different developmental stages.^27^ Another report showed temporal increase of N^6^medA levels in the mitochondrial DNA of *C. elegans*, Drosophila, and dog.^28^ Strong positive correlation between cortical N^6^medA levels and chronological age (**Figure 1C**) suggests that adenine methylation may serve as a new biomarker of human aging and could potentially be used to develop adenine methylation clocks.

Genome wide mapping of N^6^medA in the prefrontal cortex of female subjects of different ages and AD status using MeDIP-seq identified up to 350,000 N^6^medA peaks. Approximately a half of them (148,074) were shared between young female (19 Y.O.), senior female (81.8 Y.O.), and AD patient (82.5 Y.O.) samples, while the remaining peaks were shared between 2 groups only or were unique. The unique peaks found in AD patients were associated with genes involved in GABAergic synapse, glutamatergic synapse, morphine addiction, and purine metabolism (**Figure 2C)**, while adenine methylation in the age matched control was found in genes associated with regulation of actin cytoskeleton, autophagy, infection, and adherens junction (**Figure 2D**). For healthy subjects with no AD pathology, younger age was associated with adenine methylation in genes involved in cardiomyopathy, Ras signaling, and endocytosis (**Figure 2F**), while older age was associated with N^6^medA peaks in genes responsible for focal adhesion, yersinia infection, ErbB signaling, and adherens junction pathways such as axon guidance and glutamatergic synapses (**Figure 2E)**.

An alternative methodology for mapping N^6^medA (NAME-Seq) was also employed due to the high probability of false positives in MeDIP-seq. Unlike MeDIP-seq, NAME-Seq is based on nitrite conversion and polymerase extension.^41^ NAME-Seq was used to map N^6^medA across the genome of the same three samples used for MeDIP-seq (**Figure 2**) and an additional patient with mild cognitive impairment (**Table 3**). NAME-seq- revealed 26,628 and 53,198 N^6^medA peaks per sample (**Table 3**); unlike MeDIP-seq results, most of the methylated adenine peaks were unique, with only 46 peaks observed in all 4 individuals (**Figure 3A**). A large fraction of the methylated A peaks was found in introns (72-73%), followed by promoter regions (22-24 %), exons (1.5-1.8 %), and less than 1% in intergenic regions (**Figure 3D**). The most conserved sequence motif for adenine methylation was GGAAG**A**GTAG (**Figure 3B**). The majority of peaks found in AD patients are associated with focal adhesion, morphine addiction, glutamatergic synapses, these pathways show a high degree of overlap with MeDIP-seq data (**Figure 3C**).

Finally, we carried out photoaffinity proteomics using diazirine-functionalized oligodeoxynucleotide probes containing N^6^medA or dA (**Table 5**, **Fig. 5A**) and nuclear protein extracts from human neuroblastoma cells (SH-SY5Y). These experiments identified 25 proteins that have functions in mRNA processing, nucleocytoplasmic transport, DNA repair, DNA replication etc. (**Figure 5B-D**). In addition, the discovery of specific protein readers of methylated adenine such as HMGN4, TRIM28, CHAF1A merit further investigation regarding their role in regulating chromatin structure together with other known DNA epigenetic marks.

## Online Methods

### Human brain tissue samples

#### Healthy aging cohort

Postmortem frozen cortical tissues (Brodmann area 9/46) from cognitively normal individuals from different age groups were obtained through the NIH NeuroBioBank.

#### NCI-MCI-AD cohort

Frozen human dorsolateral prefrontal cortex (Brodmann area 9/46) samples from participants of Religious Orders Study (ROS) were obtained from Rush Alzheimer’s Disease Center, Chicago, IL. Details on the overall study design as well as the antemortem and postmortem neuropathological assessment procedures of ROS were reported previously.^64^ The study was approved by an Institutional Review Board of Rush University Medical Center, and all participants signed informed consent, an Anatomic Gift Act, and a repository consent to allow their tissue and data to be shared. Brain samples from 45 participants were obtained, with 16, 14, 15 samples from no cognitive impairment (NCI), mild cognitive impairment (MCI), and Alzheimer’s dementia (AD), respectively. Mean age at death, education level, post-mortem interval, and sex were matched among the three groups as closely as possible upon sample selection (**Table 1**).

### Genomic DNA extraction

10-50 mg of frozen brain tissue was disrupted via TissueRuptor II (QIAGEN) in the cell lysis solution (QIAGEN) supplemented with 1mM GSH. Proteinase K (QIAGEN) was added to a final concentration of 0.1 mg/ml and the samples were incubated overnight at room temperature with rotation. The next day, RNase A (QIAGEN) was added to a final concentration of 20 µg/ml and the samples were further incubated at room temperature for 2 hours. Then the samples were chilled on ice for 5 mins and extra proteins were precipitated by adding protein precipitation solution. At last, the DNA in the supernatant was precipitated by standard isopropanol precipitation procedure and redissolved in EB buffer supplemented with 1mM GSH.

### Genomic DNA Digestion

10 to 20 µg genome DNA was spiked with 2 fmole [^13^CD_3_ ^15^N_5_]-N^6^medA as internal standards. The resulting solution was then mixed with 25 units DNase I (recombinant, from Pichia pastoris),and incubated at room temperature overnight. After that, 2 mUnits phosphodiesterase I (type II, from Crotalus adamanteus venom) and 25 units alkaline phosphatase (recombinant, from Pichia pastoris) were added, and the samples were incubated at 37° C for another hour. The enzymes in the resulting solution were then removed with Nanosep 10K filter.

### Solid-Phase extraction

The filtered DNA samples were then completely dried down and reconstituted in 500 µl MiliQ water. 100 µl of samples is reserved for dA quantification via UV on HPLC. The remaining 400ul samples were further purified to enrich the N^6^medA fraction through solid-phase extraction (SPE) cartridge (Strata-X 33 μm, 30 mg/1 ml, Phenomenex, Torrance, CA). After sample loading, the cartridge was sequentially washed with 1ml 10% and 20% methanol and the N^6^medA fraction was eluted with 1ml 50% methanol. The eluted fraction was then completely dried down and reconstituted in 20 µl 5mM ammonium acetate.. The nanoLC/NSI-MS-MS analysis of N^6^medA was carried out on a Q-Exactive mass spectrometer (Thermo Scientific, Waltham, MA) interfaced with a Dionex Ultimate 3000 UHPLC (Thermo Fisher, Waltham,MA, USA). Chromatographic separation was achieved on a nanoLC column (0.075 mm × 150 mm) created by hand packing a commercially available fused-silica emitter (New Objective, Woburn, MA) with Luna 5 μm C18(2) 100 Å media (Phenomenex, Torrance, CA). The samples were eluted at a flow rate of 0.3 μL/min with a gradient of 5 mM ammonium acetate (A) and acetonitrile (B). Solvent composition was linearly changed from 2 to 17% B over 20 mins, then linearly increased to 70% B over the next 2 mins, and maintained at 70% B for 5 mins. The solvent composition was then returned to initial conditions (2% B) and re-equilibrated for 5 mins. The mass spectrometer was operated in a positive ion mode using a PRM experiment with FWHM of 15s. The PRM experiment were conducted at 70,000 resolution using a inclusion list of m/z 266.1248 [M+H]+ for N^6^medA and m/z 275.1312 [M+H]+ for [^13^CD_3_ ^15^N_5_]-N^6^medA and a AGC target of 1 × 10^6^, maximum IT of 50 ms, an isolation window of 0.4 m/z, a collision energy of 16. The ion fragment resulting from the neutral loss of the 2’-deoxyribose was used for quantitation. The samples information was blinded during sample preparation and LC/MS-MS analysis in order to avoid any artificial bias toward a specific group of samples.

### N^6^medA MeDIP-seq

Genomic DNA from healthy young female, healthy senior female and AD senior female’s prefrontal cortex was purified with general ethanol precipitation method. For each sample, 5 μg DNA was sheared to 150–250 bp with Covaris sonicator. Then, the DNA fragments were immunoprecipitated with N^6^medA specific antibody (5μg for each reaction, 202-003, Synaptic Systems) overnight at 4°C. The N^6^medA containing fragments were further immunoprecipitated with protein G dynabeads (Invitrogen #10004D 30mg/mL) at 4°C for 2 hours. At last, the N^6^medA containing fragments were eluted via N^6^medA nucleotide competition method and purified via ethanol precipitation. The sequencing library of these N^6^medA containing DNA fragments or input control was constructed following the instruction of NEBNext Ultra II kits (New England Biolabs). Briefly, this procedure includes end preparation, adaptor ligation, AMPureXP beads purification, PCR enrichment of adaptor ligated DNA and a final round of AMPureXP beads purification. For human samples, the libraries were then pooled and sequenced on illumina Nextseq 550 instrument in single end 75bp mode, at an average depth of 60 million reads per sample. For mouse samples, the libraries were pooled and sequenced on illumina Novaseq 6000 instrument in paired end 50bp mode, at an average depth of 60 million reads per sample.

### N^6^medA MeDIP-seq data analysis

Raw fastaq files were trimmed of low-quality bases and adapter contamination with TrimGalore! 0.4.4_dev (http://www.bioinformatics. babraham.ac.uk/projects/trim_galore/) in single-end mode using Cutadapt version 1.18 (https://journal.embnet.org/index.php/embnetjournal/article/view/200) and fastqc version 0.11.7. The filtered reads were aligned to homo sapiens GRCh38 via bowtie2^65^ version 2.3.4.1. Sam files were then converted to sorted bam files using samtools^66^ version 1.9 and duplicate reads were marked by picard-tools (https://broadinstitute.github.io/picard/) version 2.18.16. N^6^medA peaks were called via macs2 using the input-seq file as control at default settings but in –nomodel mode. Peak intersections were done via bedtools “intersection” function. Peakcount frequency profiling along genes, peak annotation were done in ChIPseeker^67^ and KEGG pathway analysis were done in ClusterProfiler.^68^ Motif discovery were conducted in STREME^69^ version 5.4.1.

### NAME-Seq methodology

One µg of genomic DNA (eluted in water) was first sonicated to around 800bp (Covavis, S220) with the following parameters: microTUBE AFA Fiber Snap-Cap, Peak Incident Power (105W), Duty Factor (5%), Cycles per Burst (200), Treatment Time (50s). Sheared DNA was denatured by incubating at 95 °C for 5 min and on ice for 5min. The denatured DNA was then treated with 1M Sodium nitrite (NaNO_2_) and 2.3% acetic acid (AcOH) at 37 °C for 2 hours. Nitrite-treated DNA was purified using Oligo Clean & Concentrator Kits (Zymo Research, D4060). The library preparation workflow after nitrite treatment was adapted from the previous post bisulfite adapter tagging method (PBAT).^30^ Briefly, to the nitrite-treated DNA solution, we added 1.25µL 10mM dNTPs (NEB, N0447S), 4µL 100µM Biotin-R2-N9 primer (5’-[Btn]CAGACGTGTGCTCTTCCGATCTNNNNNNNNN-3’), 5 µL 10X DNA polymerase buffer according to manufactory’s instructions and adjusted the final volume to 50 µL with water (Sigma-Aldrich, W4502). The mixed solution without DNA polymerase was incubated at 94 °C for 5 min and 4 °C for 5min. The program was then paused and 1.5 µL 50U/µL KF-exo^-^ (NEB, M0212M) was added to the solution. The solution was mixed and incubated at 4 °C for 15min, raised to 37 °C at a ramp rate of 0.1 °C/s, and incubated at 37 °C for 90 min to finish the first strand synthesis. Next, the enzyme activity was killed by incubating the tube at 70 °C for 10 min.

DNA after the first random priming extension was purified twice with 1.2X Ampure XP beads (Beckman, A63881) according to the manufactory’s instruction and eluted to 50 µL of 10 mM Tris-HCl, pH = 7.5 to remove the un-extended biotin primers. 20 µL Dynabeads MyOne Streptavidin C1 beads (Invitrogen, 65001) were washed and eluted in 50µL 2X B&W buffer (10 mM Tris-HCl (pH 7.5), 1 mM EDTA, and 2 M NaCl) and combined with 50 µL eluted DNA. The beads-DNA mixture was incubated at room temperature with constant rotation for 30min. The DNA bond beads were collected and washed by the following buffers: once with 2X B&W buffer, twice with fresh prepared 0.1 N NaOH (incubate for 2 min during each wash), once with 2X B&W buffer and once with 10 mM Tris-HCl, pH = 7.5.

The second random priming extension was performed on streptavidin beads by the following steps: the washed beads were suspended by 50 µL random priming buffer (1.25 µL 10 mM dNTPs (NEB, N0447S), 4 µL 100 µM R1-N9 primer (5’-CTACACGACGCTCTTCCGATCTNNNNNNNNN-3’), 5 µL 10X DNA polymerase buffer and the final volume was adjusted to 50 µL with water). The mixed solution without DNA polymerase was incubated at 94 °C for 5min and 4 °C for 5 min. The program was then paused and 1.5 µL 50U/µL KF-exo^-^ (NEB, M0212M) was added to the solution. The solution was mixed and incubated at 4 °C for 15 min, raised to 37 °C at a ramp rate of 0.1 °C/s, and incubated at 37 °C for 30min. The enzyme activity was then killed by incubating the tube at 70 °C for 10 min. Beads were collected and eluted in 50 µL *Bst-*3.0 DNA polymerase mixture (5 µL Isothermal Amplification Buffer II, 3 µL 100 mM MgSO4, 1.25 µL dNTP, 1 µL 8U/µL *Bst*-3.0 DNA polymerase (NEB, M0374S), and 39.75µL water) and incubated at 65 °C for 30min to complete the second strand synthesis.

Beads were next collected and resuspended using 50 µL Phusion elution buffer (10µL 5X Phusion HF buffer, 1.25 µL dNTP, 4 µL 10 µM R2-22nt primer (5’-CAGACGTGTGCTCTTCCGATCT-3’), 1µL 2U/ µL Phusion High-Fidelity DNA Polymerase (NEB, M0530S), and 33.75µL water). The solution was incubated at 94 °C for 5 min, 55 °C for 10 min, and 72 °C for 30 min to elute double-strand DNA from streptavidin beads. The supernatant was transferred to a new tube and treated with 1 µL Exonuclease I (NEB, M0293S) at 37 °C for 15 min and 70 °C for 10 min to remove excessive R2-22nt primer. Exonuclease I treated DNA was then purified by 1.0X Ampure XP beads to remove the short fragments. The final indexed libraries were constructed using NEBNext Multiplex Oligos for Illumina (NEB, E7335S/E7500S/E7710S/E7730S) and Q5 High-Fidelity 2X Master Mix (NEB, M0492S) (cycle number was determined by qPCR with PowerUP SYBR Green Master Mix using R1-22nt (5’-CTACACGACGCTCTTCCGATCT-3’) and R2-22nt primers (Applied Biosystems, A25742)). To increase the efficiency for identifying N^6^medA, NAME-Seq could be coupled with MeDIP (DIP-NAME-Seq), which is employed in this study. Samples were sequenced using Illumina NovaSeq 6000.

### NAME-seq Data Analysis

Analysis workflow is similar to the previously reported NT-seq.^17^ The scripts used for analysis are available at (https://github.com/TaoLabBCM/NAME-seq). Briefly, sequencing reads were deduplicated using BBMAP (v38.84)^32^ Clumpify package and then trimmed using Cutadapt (v1.18)^33^. To align all nitrite treatment converted reads (A to G and C to T mutations) to reference genome, FASTQ reads and FASTA reference were converted to AT-only format (convert all purine (A/G) to A and all pyrimidine (C/T) to T). Converted FASTQ reads were aligned to converted reference using Bowtie2 (v2.3.5.1)^34^. AT only SAM files were converted back to SAM files with original reads using custom python scripts. NM (Number of mismatches between the sequence and reference) and MD (String encoding mismatched reference bases) tags in SAM files were recalculated using Samtools (1.9)^35^ calmd command to obtain the mutation pattern of original reads. The alignments in SAM files were further filtered by recalculated NM and MD tags to remove unconverted reads and reads with unwanted base mutation (number of mismatch >= 2, number of mismatches at A or C position/total number of mismatches >= 0.8, and no mismatch at T position). The base count at each genomic location was generated by Igvtools (v2.5.3)^36^ count command using filtered SAM files, and the base count files were used to calculate A to G frequency and A to T frequency at adenine positions and the C to T frequency at cytosint positions using custom python scripts. 6mA motifs in K562 were identified using MEME-suite (v5.4.1)^37^.

### N^6^medA affinity proteomics

#### Nuclear lysate preparation

Human-derived neuroblastoma (SH-SY5Y) cells were cultured in 50/50 EMEM/F-12 (Gibco) media supplemented with 10% FBS and 1x PS. The cells were harvested via trypsin digestion, washed twice in cold PBS and then resuspended in cell lysis buffer (10 mM Tris pH = 8, 10 mM KCl, 10 mM MgCl_2_, 10 mM DTT) supplemented with 1x complete EDTA-free protease inhibitor cocktail tablets (Roche Diagnostics GmbH) and 0.5% phosphatase inhibitor cocktail 2 (Sigma). The cells were homogenized with 5 passes through a syringe with a 25 gauge needle and then centrifuge at 2600 rpm for 8 mins. After that, the pellet was resuspended in nuclear extraction buffer (10 mM Tris pH = 7.4, 10 mM KCl, 10 mM MgCl_2_, 10 mM DTT, 250 mM NaCl) at half the cell lysis buffer volume, supplemented with 1x complete EDTA-free protease inhibitor cocktail tablets and 0.5% phosphatase inhibitor cocktail 2, and incubated on ice for 1 h. Then the samples were centrifuged at 14,800 rpm at 4 °C for 30 min. The pellet was discarded, 10% glycerol was added to the supernatant, and the nuclear protein lysate was stored at -80 °C until use.

#### DNA oligonucleotide preparation

DNA oligomers were synthesized in-house on a MerMade 4 DNA Synthesizer. 5-aminoallyl-dU CEP was purchased from BOC Sciences; all other reagents used for DNA synthesis were purchased from Glen Research. The forward, biotinylated strand was synthesized with a single 5-aminoallyl-dU, purified using HPLC and desalted using NAP™-5 ColumnsSephadex™ (Cytiva). The biotinylated strand (160 µM) was then reacted with 2,5-dioxopyrrolidin-1-yl 3-(3-methyl-3H-diazirin-3-yl)propanoate in DMSO (10 mM) (synthesized in-house) and Na-HEPES (100 mM) (Sigma), pH 7.5 for 3 hours in the dark at room temperature. The biotinylated-diazirine oligomer was then purified via HPLC and again desalted using NAP™-5 ColumnsSephadex™.

Two complimentary strands containing either adenosine (control) or N6-methyladenosine (modified) were synthesized in-house on a MerMade 4 DNA Synthesizer. Each complimentary strand was annealed to the biotinylated-diazirine oligomer by adding equimolar amounts to 1X DNA annealing buffer (10 mM Tris-HCl, 50 mM NaCl, pH 8.0), heating to 94 °C for 5 minutes, and then letting the mixture cool to room temperature.

### Affinity pulldown and on-bead digestion

For each sample, 400 pmol of the biotinylated dsDNA bait (either control or modified; 3 replicates of each) was incubated with 500 µg of nuclear extract, diluting to a final volume of 500 µL with Buffer C (20 mM Tris-Cl pH 7.5, 50 mM NaCl, 1 mM EDTA, 1 mM DTT, 20% glycerol), by rotating at 4 °C for 4 hours in the dark. The mixture was exposed to 365 nm light for 10 minutes to activate the photolinker and crosslink the dsDNA to affinity-bound proteins. Next, the mixture was added to 70 µL of Streptavidin Mag Sepharose™ (Cytiva) that had been washed 3 times with Tris-buffered saline (50 mM Tris-HCl pH 7.5, 150 mM NaCl) and once with Buffer C. To immobilize the DNA-protein crosslinks (DpC) on the beads, the bead mixture was rotated overnight in the dark at room temperature. The DpC-decorated beads were washed 3 times with Buffer C and the supernatant was discarded.

The samples were resuspended in 100 µL reduction buffer (50 mM NH_4_HCO_3_, 2mM TCEP) and incubated at 65 °C for 1 hour. Then 1 µL alkylation buffer (100 mM iodoacetamide in 50 mM NH_4_HCO_3_) was added to each sample and incubated for 30 mins at room temperature in the dark, after which the supernatant was discarded. The samples were resuspended in 200 µL digestion buffer (50 mM NH_4_HCO_3_, 500 mM urea, 7.5 ng/µL trypsin, pH 8) and incubated overnight at 37 °C. The supernatant was kept and the beads were washed once with 200 µL PBS and 200 µL HPLC-grade water; both washes were retained and added to the supernatant. The combined supernatant was dried via vacuum centrifugation. Tryptic peptides were desalted using Pierce™ C18 spin columns (ThermoFisher Scientific) per manufacturer’s guidelines with LCMS-grade acetonitrile, water, and TFA. Peptides were dried using vacuum centrifugation and reconstituted in 10 µL LCMS-grade water containing 0.1% formic acid.

#### LC-MS/MS analysis

The six peptide samples were analyzed on an Orbitrap Exploris 480 mass spectrometer (ThermoFisher Scientific, Waltham, MA) interfaced with a Vanquish UHPLC (ThermoFisher Scientific, Waltham,MA, USA). The UHPLC was run in nanoflow mode with a reverse-phase nanoLC column (0.075 mm × 150 mm) created by hand packing a commercially available fused-silica emitter (New Objective, Woburn, MA) with Luna 5 μm C18(2) 100 Å media (Phenomenex, Torrance, CA). The samples were eluted at a flow rate of 0.3 μL/min with a gradient of water (A) and 95% acetonitrile (B). Solvent composition was linearly changed from 5 to 40% B over 45 mins, then linearly increased to 95% B over the next 5 mins, and maintained at 95% B for 5 mins. The solvent composition was then returned to initial conditions (5% B) and re-equilibrated with 5 times the column void volume. The mass spectrometer was operated in positive mode using a Full MS/dd-MS2 experiment with an expected LC peak width of 10 s. In the full scan mode, resolution was 60,000 with an AGC target of 1 × 10^6^, a maximum IT of 40 ms, and a scan range of 350 to 1200 m/z, an isolation window of 1.6 m/z, and dynamic exclusion time of 30 s, and a normalized collision energy of 30.

#### N^6^medA affinity proteomics data analysis

Raw mass spectrometry files were analyzed using FragPipe computational platform (v22.0) (). The spectral identification was performed using MSFragger v4.1 against the UnitProt Human reference proteome (UP000005640). Search parameters included Trypsin/LysC specificity, precursor/fragment mass tolerances of 20 ppm, and fixed carbamidomethylation. Peptide−spectrum matches were validated using Percolator and filtered with a False Discovery Rate (FDR) cutoff of 0.01. The quantification of the data set utilized Label-Free Quantification (LFQ) along with IonQuant v1.10.27 with Match-Between-Runs (MBR) enabled. MaxLFQ normalization was applied to corrected for sample loading differences. The statistical analysis was performed using R. Any missing values were categorized as Missing Not At Random (MNAR) and imputed using a left-censored normal distribution approach. Significantly enriched proteins were tested using two-sided Student’s T-tests and the *P*-values were corrected using the Benjamini-Hochberg (BH) method. KEGG pathway analysis was performed on the complete data set, ignoring the p-values to obtain broad, system-wide function shifts.

## Reporting summary

Further information on research design is available in the Nature Portfolio Reporting Summary Linked to this article.

## Data availability

All data generated or analyzed during this study are included in this published article and its supplementary information file. Additional information is available from the corresponding author upon reasonable request.

## Acknowledgements

We thank David Bennett, Debra Magnuson, Karen Skish, and Gregory Klein at Rush University Alzheimer’s Research Center and staff members at NIH NeuroBioBank for selecting and providing the postmortem human brain specimens. This work was supported in part by funds from the University of Minnesota Academic Health Center, Masonic Cancer Center, UMN College of Pharmacy, and the Department of Medicinal Chemistry. L.L. and A.J. were supported by the grant RF1AG056976 from the National Institute on Aging of the National Institutes of Health.

## Ethics declarations

