## Supplementary Information for "DNA Adenine Methylation Clock in Brain Aging and Alzheimer’s Disease Progression"

**Figure S1:** NAME-seq results for $N^6$medA in genomic DNA isolated from prefrontal cortex of 3 human subjects: a young female (19 y.o.), a senior female control (81.8 y.o), a female with mild cognitive impairment (MCI, 82.9 y.o.), and a female AD patient (82.5 y.o). **a-d,** NAME-seq results for GO enrichment analysis on genes containing $N^6$medA peaks in AD female, senior healthy female and young healthy female respectively. **e-f,** Top sequence motif for adenine methylation in mild cognitive impairment patient (MCI) and senior healthy respectively.

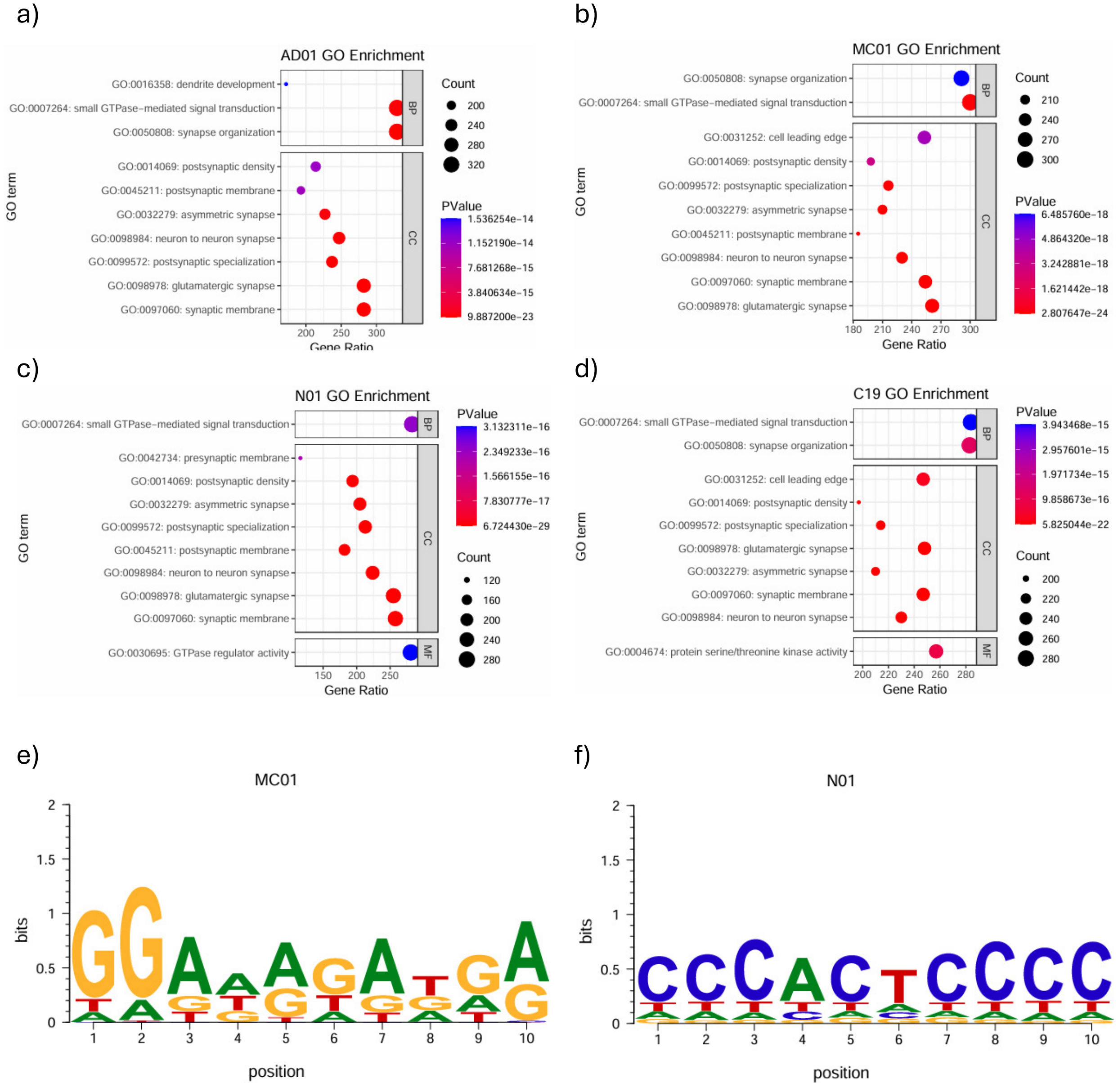

**Figure S2a-c:** Cross validation of N$^6$medA peaks using MeDIP-seq and NAME seq methodologies in genomic DNA isolated from prefrontal cortex of 3 human subjects: a female AD patient (82.5 y.o), a senior female control (81.8 y.o), and a young female (19 y.o.). Approximately 10% of N$^6$medA sites were identified in NAME seq were cross valided by MeDIP seq.

a)

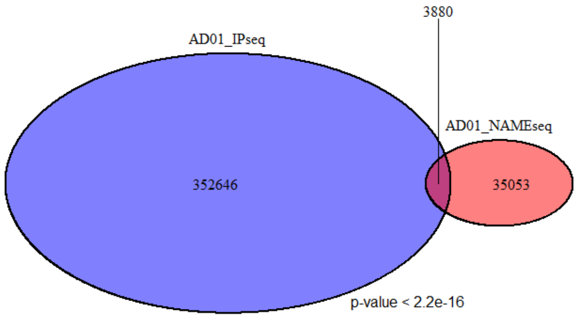


b)

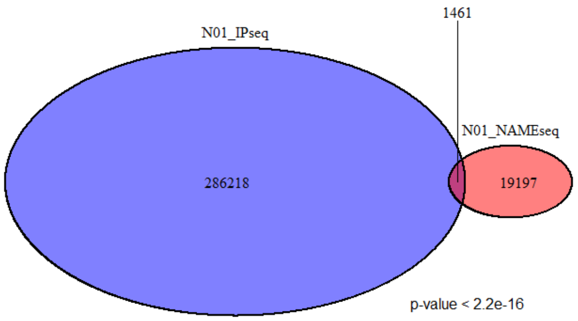


c)

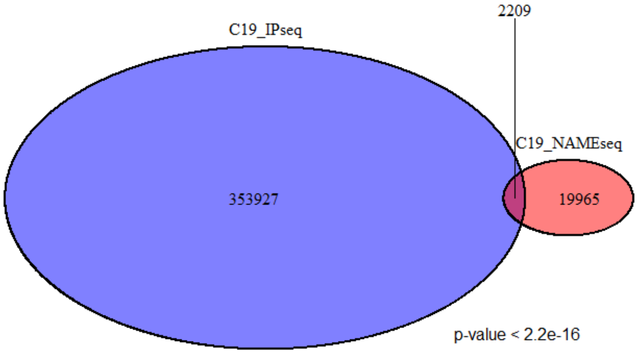

**Figure S3: Integration of MeDIP-seq and NAME-seq results for $N^6$medA in genomic DNA isolated from prefrontal cortex of 3 human subjects: a young female (19 y.o.), a senior female control (81.8 y.o), a female with mild cognitive impairment (MCI, 82.9 y.o.), and a female AD patient (82.5 y.o).**

**a-d,** integration results for GO enrichment analysis on genes containing $N^6$medA peaks in AD female, senior healthy female and young healthy female respectively. **e-f,** integration results for top sequence motif for adenine methylation in senior healthy female and young healthy female respectively.

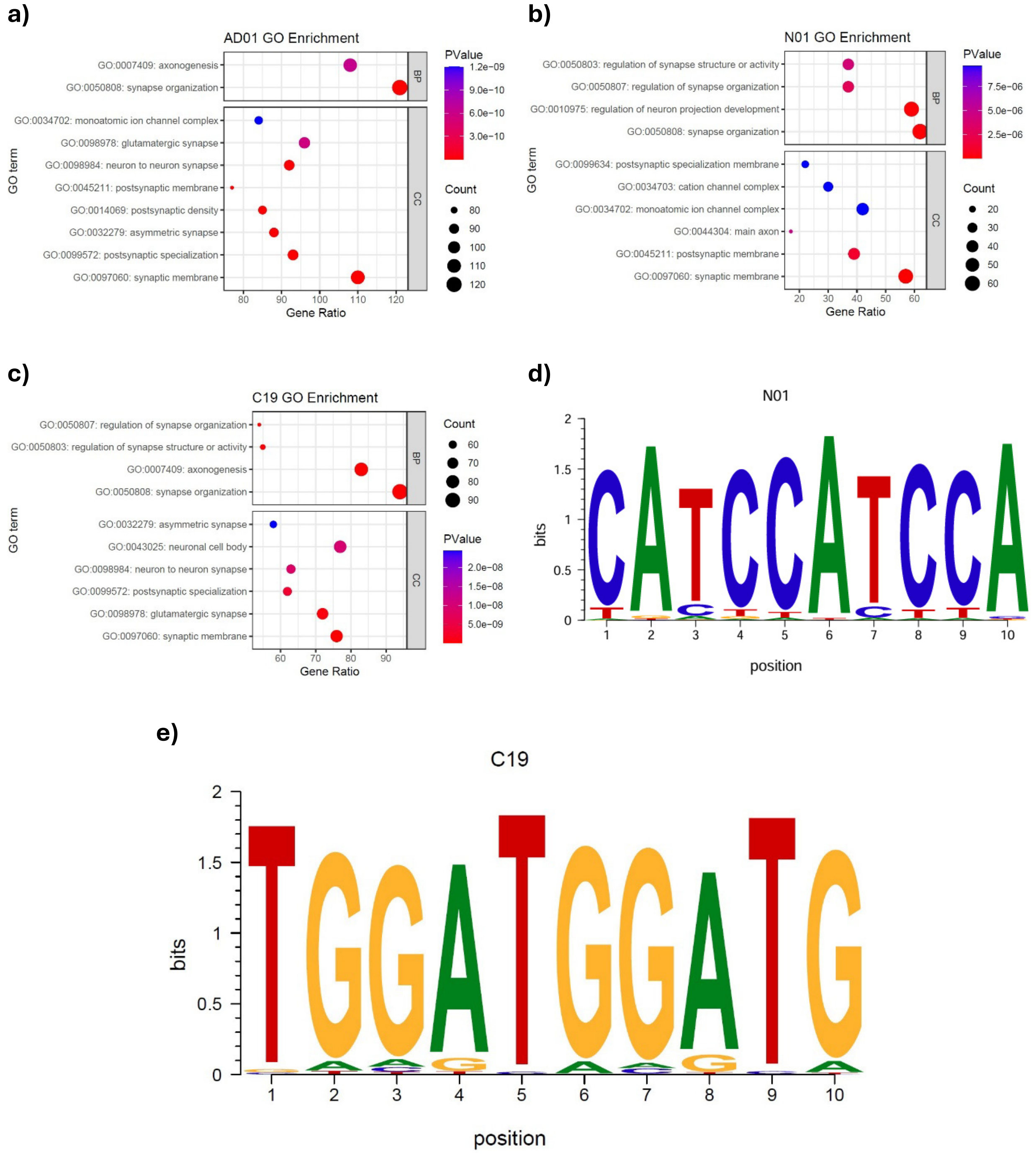

**Figure S4: Characterization of protein readers of N$^6$medA via affinity proteomics for DNA repair pathway. a,** volcano plot showing the quantitative analysis of proteins identified in DNA repair pathway. **b,** Hierarchical clustering heatmap showing top 25 protein abundance identified in DNA repair pathway upon N$^6$medA affinity pulldown in SH-SY5Y cells nuclear extract.

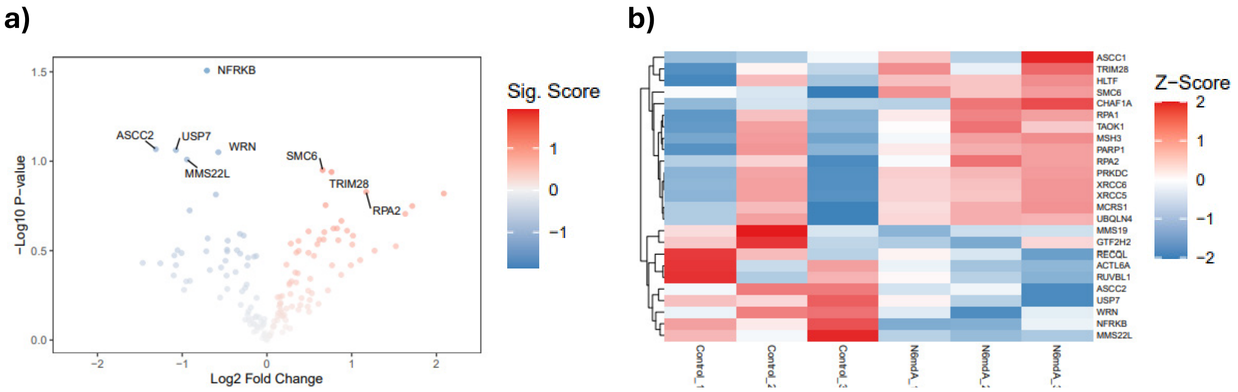


**Table S1:** Number of N$^6$medA sites per cell calculated using different methods

| Method | N$^6$medA sites per cell |
|---|---|
| LC-MS | ~5700 |
| NAME-seq | ~30000 |
| MeDIP-seq | ~300000 |
| Overlap | ~3000 |
